# Multisensory Configural Learning of a Home Refuge in the Whip Spider *Phrynus marginemaculatus*

**DOI:** 10.1101/2020.09.29.318659

**Authors:** Kaylyn AS Flanigan, Daniel D Wiegmann, Eileen A Hebets, Verner P Bingman

## Abstract

Whip spiders (Amblypygi) reside in structurally complex habitats and are nocturnally active yet display notable navigational abilities. From the theory that uncertainty in sensory inputs should promote multisensory representations to guide behavior, we hypothesized that their navigation is supported by a configural, multisensory representation of navigational inputs, an ability documented in a few insects and never reported in arachnids. We trained *Phrynus marginemaculatus* to recognize a home shelter characterized by both discriminative olfactory and tactile stimuli. In tests, subjects readily discriminated between shelters based on the paired stimuli. However, subjects failed to recognize the shelter in tests with either of the component stimuli alone. This result is consistent with the hypothesis that the terminal phase of their navigational behavior, shelter recognition, can be supported by the integration of multisensory stimuli as a configural representation. We hypothesize that multisensory configural learning occurs in the whip spiders’ extraordinarily large mushroom bodies, which may functionally resemble the hippocampus of vertebrates.

## INTRODUCTION

Many arthropods are superb navigators (Menzel et al., 2005; Wehner, 2003; Cheng et al., 2009; Layne et al., 2003). Those species that inhabit structurally uncluttered environments (e.g., deserts) typically rely on sensory inputs that are organized in parallel as independent, sometimes weighted inputs (Collett et al., 2013; Müller & Wehner, 2007; Wehner et al., 2016). In environments where sensory inputs are associated with uncertainty, such as in structurally cluttered environments (e.g., dense forests), behavior guided by multisensory inputs may be advantageous (Munoz & Blumstein, 2012). Multisensory inputs, when used, can be processed independently (Wehner et al., 2016) or integrated in the form of a configural representation (Pearce, 2002; Swinton et al., 2019; Kagan & Lukowiak, 2019). In the context of navigation, a multisensory representation should reduce spatial uncertainty, and if such a representation were configural, it could reduce uncertainty even further, as error prone input from one cue would have little control over behavior when not processed with the companion, configural input(s). Nonetheless, reports of configural, multisensory control of arthropod navigation are scarce. In insects, integration of celestial and wind inputs interact to support the directional movements of dung beetles (Dacke et al., 2019). In ants, true multisensory-configural representations binding visual and odor inputs can guide locating a nest entrance (Steck et al., 2012) or a feeder (Buehlmann et al., 2020). Beyond the context of navigation, the ant *Lasius niger*,can learn the conditional relationship between an odor and color cue to locate food warrants (De Agrò et al., 2020).

Multisensory integration in arachnid communication is well documented (Hebets, 2005; Higham & Hebets, 2013), but we are unaware of evidence that arachnids can form configural representations of different sensory modalities to guide navigation. Nocturnally active whip spiders of the order Amblypygi are excellent navigators, successfully returning to their home refuge after a night of hunting and even after experimental displacement (Beck & Görke, 1974; Hebets et al., 2014; Bingman et al., 2017). In the Florida Keys, the whip spider *Phrynus marginemaculatus* resides in dense scrub habitat cluttered with limestone and plant litter. Wiegmann et al. (2016) proposed that, in such an environment, the configural learning of multisensory inputs facilitates whip spider navigation. Here, we hypothesized that *P. marginemaculatus* can learn a configural, multisensory representation of a shelter location, which if supported, would demonstrate for the first time in arachnids that configural, multisensory representations can guide navigation behavior.

## MATERIALS AND METHODS

### Subjects

Twenty one *P. marginemaculatus* (C.L. Koch 1840), collected in Florida (Key Deer National Wildlife Refuge, Big Pine Key, Monroe County, FL) and lab-borne progeny of subjects collected in Florida, served as the experimental subjects. The sex of the subjects was not recorded. We trained and tested six naïve *P. marginemaculatus* in an olfactory-only shelter-discrimination task. In a separate experiment, we trained and tested three naïve animals and three subjects from the olfactory-only task in a tactile-only shelter-discrimination task. The three subjects tested on both tasks experienced the tactile-only task second, which was separated by at least four weeks from the olfactory-only discrimination task to minimize any cue interference. (The performance of the three subjects tested on the tactile-only discrimination task after being trained on the olfactory task was indistinguishable from the three subjects tested solely on the tactile-only task.) In the last experiment, we trained twelve naïve animals on the critical multisensory shelter-discrimination task.

### Single-cue shelter discrimination

Earlier studies with experimental designs that differed from ours demonstrated that *P. marginemaculatus* can locate a shelter using olfactory or tactile cues (Wiegmann et al., 2019; Santer & Hebets, 2009). Here, we trained and tested subjects on olfactory-only (N = 6) and tactile-only (N = 6) shelter discrimination tasks under conditions similar to those used in the primary multisensory discrimination experiment to better compare performance of single-cue and multi-cue guided behavior in the experiments of the current study.

Olfactory-only discrimination training occurred over four consecutive nights (Figure 1). A subject was placed in a pre-exposure shelter made of opaque PVC pipe, which contained 10μL of geraniol (C_10_H_18_O, Sigma-Aldrich, Product number 163333; three subjects) or 1-hexanol (C_6_H_14_0, Sigma-Aldrich, Product number 471402; three subjects) two hours before lights-out occurred. *P. marginemaculatus* can use either odor to locate a shelter (Wiegmann et al., 2019). Pre-exposure served to facilitate the association of an odor with the safety of the shelter. After lights out the subject was transferred to a training arena containing two shelters identical in size to the pre-exposure shelter. One shelter, which was accessible, contained the pre-exposure odor (CS+). The second shelter, whose entrance was blocked by mesh, contained the other, unconditioned odor (CS−). After five-minutes, we allowed subjects to leave the pre-exposure shelter and the shelter was removed. (If, after five minutes, the subject failed to exit, we gently coerced it out with a paintbrush.) The next morning (lights on), the subject was enclosed in the CS+ shelter, where it was held until testing. (If it was not in the shelter, we gently coerced it inside.)

**Figure 1.**
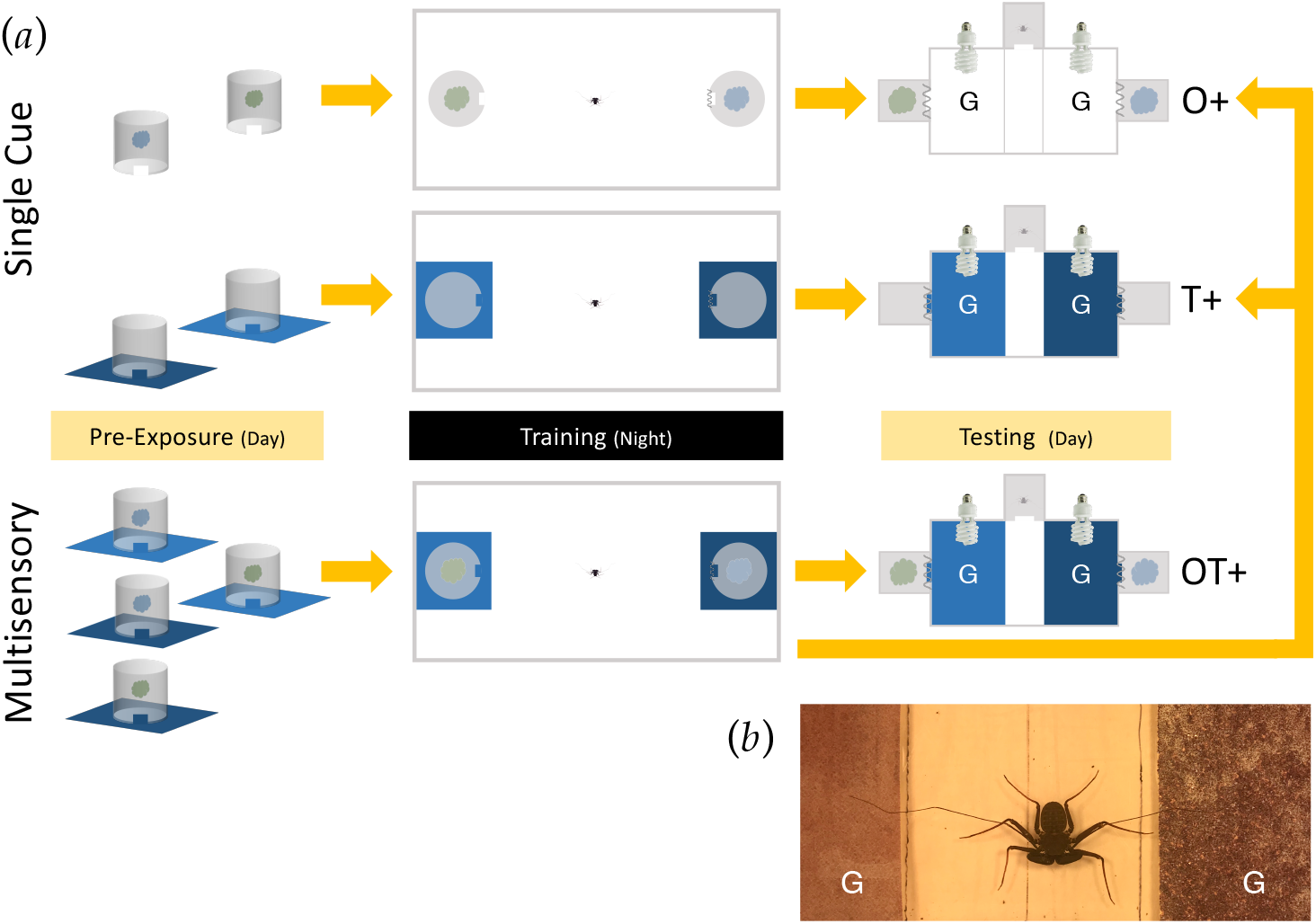
(a) Schematic of training and tests. Subjects were placed in a pre-exposure shelter (diameter, 10.2 cm; height, 12 cm) that contained a CS+ odor (blue and green “clouds”) or sandpaper (light and dark blue surfaces) for the single cue experiment and a CS+ paired odor and sandpaper (four combinations) for the multisensory experiment. At night, we released subjects into a training arena (length, 1.0 m; width, 0.5 m; height, 30.0 cm) that contained two shelters. One shelter, which was accessible, contained the CS+. The other shelter, which was inaccessible, contained the CS−. (The sandpaper extended beyond the boundaries of the shelters.) The next morning, subjects were tested. The floor of the test arena (length, 29.0 cm; width, 14.0 cm; height, 9.5 cm) was demarcated into two goal areas, G (length, 14.0 cm; width, 10.5 cm). We separated goal areas by a central “neutral zone” (length, 14.0 cm; width, 7.0 cm). A “start chamber” (length, 7.0 cm; width, 5.0 cm, height, 2.5 cm) opened into neutral zone. Two shelters at the ends of the arena, which could *not* be entered because of a mesh barrier, contained the CS+ and CS− odors. The CS+ and CS− sandpaper occupied their respective goal areas. We illuminated the test arena by two light sources. O, olfactory; T, tactile. See Methods for more details. (b) Image of a subject in the neutral zone during a test. Note the antenniform legs could be extended to touch both sandpaper surfaces. The average prosoma width of the test subjects was approximately 0.9 cm and the antenniform legs, when extended, were approximately 9.0 cm.

We conducted tests two to four hours after lights-on (Figure 1). Each end of the test arena had an entrance to a shelter, one of which contained 10μL of the CS+ odor (side determined randomly) and one that contained 10μL of the CS− odor. A mesh door rendered both shelters inaccessible. The arena was illuminated by two 13-watt Phillips Mini Twister (120V, 60 Hz) light bulbs to motivate subjects to escape the light and locate a shelter. Test trials lasted 10 minutes. We recorded the time a subject spent in goal areas near shelter entrances. Our dependent measure to assess the strength of the learned association to the CS+ shelter was the “association index” (θ), which was the time spent near the C+ shelter divided by the total time spent near both shelters. After testing, we placed individuals in a pre-exposure shelter until lights out, when we again placed them in the training arena. This sequence was repeated four times for a total of four tests per individual.

Procedures for the tactile-only discrimination task were the same, except for the conditioned stimuli (Figure 1). Three animals were trained to coarse sandpaper as the CS+ and three were trained to smooth sandpaper (3M Pro Grade Precision Sandpaper, P60 and P320 respectively). We covered the floor of the pre-exposure shelter with the CS+. In training, we placed sheets of sandpaper (28 cm X 23.8 cm) inside and around each shelter. The tactile features covered the goal areas in the test arena.

### Multisensory shelter discrimination

We trained naïve (N = 12) subjects in a multisensory discrimination task with the same olfactory and tactile cues, in this situation paired, where we balanced the four possible CS+ combinations across subjects. Tests involved either the paired cues or the component stimuli alone (Figure 1). Subjects were required to meet a recognition criterion θ ≥ 0.70 with the paired stimuli on three consecutive tests before formal tests were initiated. All twelve subjects rapidly reached criterion (none were excluded from the experiment for failure to reach criterion). Testing involved four blocks of three tests, where each block included an odor-only, tactile-only and a paired-stimuli test. Test order within blocks was determined randomly except for the first test in the first block, which was restricted to a single-cue test to avoid habituation after criterion tests. We conducted tests over 12 days, one test per day. Procedures otherwise followed those described for the single-cue experiment.

### Quantitative analysis and statistical procedures

Larger values of the association index (0 ≤ θ ≤ 1) indicate a stronger conditioned association between the CS+ and access to shelter. For the single-cue experiment, indices from the four tests for each subject were averaged and one sample *t*-tests were used to compare mean indices to the random expectation of θ = 0.5. For the multisensory experiment, the means for each individual within test-trial types were averaged and likewise compared to θ = 0.5. In addition, a repeated measures ANOVA, with Tukey HSD *q* post-hoc tests to control for Type I error, was used to compare performance between the single-cue and paired-cue tests. Lastly, we compared performance in single-cue tests of the multisensory experiment to performance in the single-cue experiments with two sample *t* tests.

## RESULTS

We provide goal-area occupancy times for all subject-test trials in Supplementary Table 1.

### Single cue shelter discrimination

The mean association index for the odor-only shelter discrimination task was 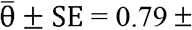 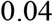 and differed significantly from the chance expectation θ = 0.5 (Figure 2, *t*_5_ = 7.67, *p* < 0.0006). The mean index for the tactile-only shelter discrimination task was 0.69 ± 0.04 and likewise differed from chance (Figure 2, *t*_5_ = 5.09, *p* = 0.0038). Hence, in the experimental context of this study, *P. marginemaculatus* learned to recognize an accessible shelter cued solely by an odor or tactile stimulus (see also Higham & Hebets, 2013; Hebets et al., 2017; Wiegmann et al., 2019).

**Figure 2.**
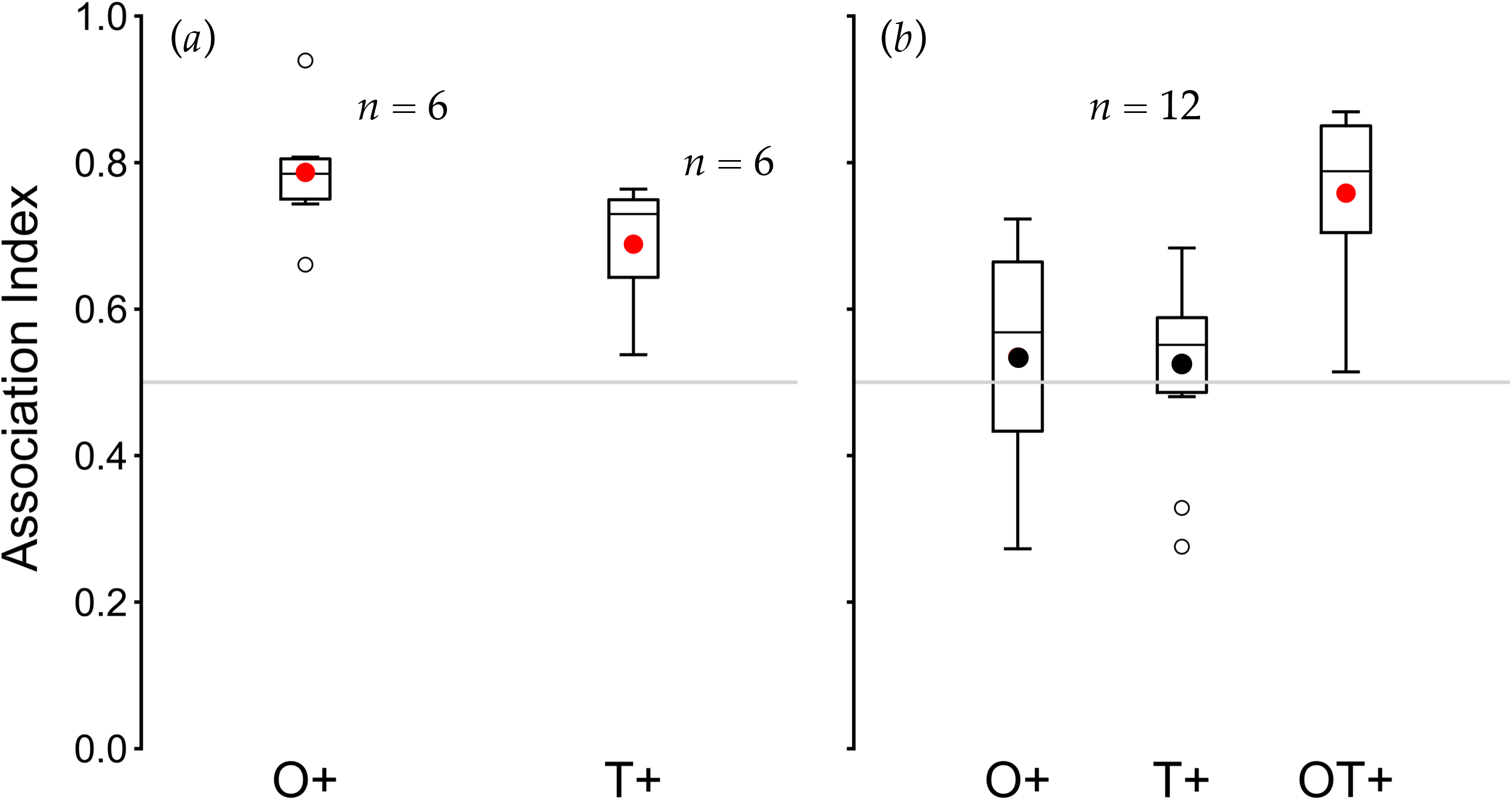
Box plots of association indexes in tests for (a) single cue and (b) multisensory shelter discrimination experiments. Filled circles are group means and open circles are outliers. Lines within the standard error boxes are medians. O+, olfactory-only tests; T+ tactile-only tests; OT+, tests with paired olfactory and tactile stimuli. Index means near one indicate a preference for CS+. Red index means differ significantly the chance expectation (grey horizontal line) of θ = 0.5.

### Multisensory shelter discrimination

In the multisensory discrimination experiment subjects required on average 3.33 ± 0.19 nights to reach criterion to move on to formal test trials. The repeated measures ANOVA revealed that association indices differed across the three test conditions (*F*_2,22_ = 15.35, *p* < 0.0001). Post-hoc tests revealed that the mean index in odor-only (0.53 ± 0.05) and tactile-only (0.53 ± 0.04) tests did not differ (*q*_22_ = 0.19, *p* = 0.9804) and that performance in paired-stimuli tests (0.76 ± 0.03) was better than in either single-stimulus test (paired versus odor-only: *q*_22_ = 4.70, *p* = 0.0003; paired versus tactile-only: *q*_22_ = 4.89, *p* = 0.0002). Performance in odor-only and tactile-only tests did not differ from chance (Figure 2, *t*_11_ = 0.76, *p* = 0.4631 and *t*_11_= 0.72, *p* = 0.4876, respectively). By contrast, performance on the multisensory shelter discrimination was significantly biased toward the CS+ (Figure 2, *t*_11_ = 7.85, *p* < 0.0001).

Discrimination performance in single-cue tests in the multisensory experiment was also poorer than in analogous tests in the single-cue experiments (olfactory-only tests: *t*_16_ = 3.63, *p* = 0.0022; tactile-only tests: *t*_16_ = 2.89, *p* = 0.0105). Performance in paired-stimuli tests, however, was no better than that of subjects trained on either of the element stimuli in the single-cue experiments (olfactory cue: *t*_16_ = 0.53, *p* = 0.6054; tactile cue: *t*_16_ = 1.30, *p* = 0.2114). Together, these results indicate that the component stimulus from either sensory modality could control shelter recognition when stimuli were individually conditioned, but neither stimulus alone could support shelter recognition when we trained subjects to the paired olfactory and tactile stimuli.

## DISCUSSION

The results of this study support our hypotheses that whip spiders can use multisensory integration in support of the terminal phase of navigation, and more interestingly, construct a configural representation of a shelter; when subjects were trained to recognize a refuge characterized by both olfactory and tactile stimuli neither component stimulus by itself supported shelter recognition. As such, whip spiders are among a growing list of invertebrates shown to be capable of configural learning (e.g., the snail, Lymnaea; Swinton et al., 2019; Kagan & Lukowiak, 2019). This exclusion of behavioral control by either component stimulus is a defining feature of configural learning as described in vertebrates (Pearce, 2002). In the context of arthropod navigation, configural learning has been observed to control aspects of navigation in two species of ants (Steck et al., 2011; Buehlmann et al., 2020), but to the best of our knowledge it has not been previously reported to control arachnid navigation. Following from the signal uncertainty (Munoz & Blumstein, 2012) likely associated with the structural complexity and nocturnal habits of whip spiders, it is perhaps not surprising that they are capable of configural learning to support navigation (see Wiegmann et al., 2016).

We also expected that discrimination (correct choice performance) would be better when a shelter was associated with multiple cues compared to a single stimulus. This was not found as the performance of subjects trained and tested on paired-stimuli was no better than that of subjects trained on either of the element stimuli. However, the advantage of configural, multisensory learning has been theorized to manifest in settings where there is uncertainty in the information carried by stimuli. In retrospect, therefore, this result is perhaps no surprise as the controlled experimental conditions generated no environmental uncertainty with regard to shelter identity and, hence, no advantage to configural, multisensory learning with regard to shelter recognition.

Multisensory, configural learning in an arthropod raises the question of the underlying neural architecture. We proposed (Wiegmann et al., 2016) that the mushroom bodies, which support learning and memory in insects (Menzel et al., 2006), are the site of multimodal, sensory integration in whip spiders. Here, two observations are noteworthy. First, the mushroom bodies are exceptionally large and elaborately folded in whip spiders, suggesting their importance in the control of complex behavior and memory-based navigation (Strausfeld et al., 2009). Second, honeybees trained to discriminate a reward based on a configural representation of two odors were unable to carry out the discrimination when the mushroom bodies were inactivated (Devaud et al., 2015). Consistent with this last observation are the results from a recent electrophysiological study revealing that many mushroom body output neurons in the honeybee, *Apis mellifera carnica*, respond to multimodal inputs suggesting a role in multimodal integration and perhaps configural learning (Strube-Bloss & Rössler, 2018).

Finally, Strausfeld et al. (2009) proposed that the mushroom bodies of arthropods are functionally equivalent to the vertebrate hippocampus. Indeed, that the hippocampus appears to support configural learning in mammals (Sutherland & Rudy, 1989), although its precise role is still uncertain (Whishaw & Tomie, 1991). If the configural representation of a shelter in whip spiders were a result of multisensory integration in the mushroom bodies, it would provide support for Strausfeld et al.’s idea of some functional equivalence between the mushroom bodies of arthropods and the vertebrate hippocampus.

## Acknowledgements

We are grateful for financial support from the National Geographic Society and the National Science Foundation (IOS 1457304).

## Author contributions

KASF contributed to the experimental design, carried out data collection and analyses and the writing of the paper. DDW contributed to the experimental design, data analyses and the writing of the paper. EAH contributed to the experimental design and writing of the paper. VPB contributed to the experimental design, data analyses and writing of the paper.

